# Novel Competition test for food rewards reveals stable dominance status in adult male rats

**DOI:** 10.1101/2020.09.24.312033

**Authors:** Diana F Costa, Marta A Moita, Cristina Márquez

## Abstract

Social hierarchy is a potent modulator of behavior, that is typically established through overt agonistic interactions between individuals in the group. Once established, social ranks are maintained through subtler interactions allowing the redirection of energy away from agonistic interactions towards other needs. The available tasks for assessing social rank in rats allow the study of the mechanisms by which social hierarches are formed in early phases but fail to assess the maintenance of established hierarchies between stable pairs of animals, which might rely on distinct neurobiological mechanisms. Here we present and validate a novel trial-based dominancy assay, the modified Food Competition test, where established social hierarchies can be identified in the home cage of non-food deprived pairs of male rats. In this task, we introduce a small conflict in the home cage, where access to a new feeder containing palatable pellets can only be gained by one animal at a time. We found that this subtle conflict triggered asymmetric social interactions and resulted in higher consumption of food by one of the animals in the pair, which reliably predicted hierarchy in other tests. Our findings reveal stable dominance status in pair-housed rats and provide a novel tool for the evaluation of established social hierarchies, the modified Food Competition test, that is robust and easy to implement.

## Introduction

Social hierarchy is a multidimensional trait that has a profound impact on emotion and cognition, not only for humans ^1, 2^ but also other social species (see ^3^ for review), having important consequences for social organization, survival, reproductive success, and health of animals in a group ^4^. Indeed, adapting behavioral responses based on the social status of the interacting partner can be cost-effective and, in some cases, a crucial survival strategy. The most established view is that social hierarchy is built upon aggressive interactions ^5^, and serve as a mechanism of resource management and minimization of energy expenditure by groups of animals: once a hierarchy is established, priority access to resources is organized allowing the reduction of aggressive levels between the interacting animals ^6^. Following this view, the behavioral paradigms available for measuring social hierarchy in laboratory animals are based in the nature of agonistic interactions while defending access to resources, whether a sexual partner, food or water when they are scarce, or the defense of a territory (see ^3^ for review).

Of note, most recent advances on the identification of the neural circuits underlying the establishment of social hierarchies have been performed in mice, as a reflection of a general tendency in the field which favors the use of this species due to the exceptional genetic tools available ^7, 8^. However, important contributions have been also performed using rats ^9–11^, and importantly, Norway rats live in complex social groups in the wild. This, together with the fact of being a model system amenable to monitoring, mapping and perturbation of neuronal circuits, has motivated a wave of recent laboratory studies uncovering the diversity and sophistication of rat’s social skills ^12^. Regarding their social status, the visible burrow system (VBS) has been widely used to study the formation of hierarchies in large groups of animals, where mixed-sex rat groups living in a complex environment compete chronically for territory and resources ^13^. The VBS generates very rich behavioral data sets but is difficult to implement in most laboratories, hence, other behavioral tasks are commonly used, where animals compete for food or water under deprivation states ^14–17^. Typically, social isolation of variable durations is performed prior to testing as a means to increase territoriality, favoring strong agonistic interactions during the establishment of new hierarchies. Therefore, these tasks evaluate how a new hierarchy is established between pairs of unfamiliar, frequently isolated animals, in neutral arenas where subjects display very evident agonistic behaviors to establish dominance.

However, there are no tools available enabling to assess already established hierarchies. Focusing on the early establishment of a hierarchy is neglecting a very important and rich part of this type of social interactions: how are they maintained in stable conditions. The establishment of social hierarchy might not rely on the same mechanisms as the expression of dominance when a hierarchy is already established, and recent reports in mice indicate that this might be indeed the case ^18, 19^. However, to our knowledge, the study of the possible differences between de novo and already established social hierarchies in rats has been virtually unexplored. To this end new behavioral paradigms that evaluate social status of animals living in stable dyads are urgently needed. Preferably, the evaluation of the hierarchy would avoid manipulations that could induce long-term alterations in internal state (i.e. isolation, deprivation) that, in turn, could affect how social interactions of interest are displayed. This is a challenge, as once hierarchies are established, agonistic behaviors are minimized and the opportunities to observe dominance interactions are reduced and likely subtle.

With this aim, we developed the modified Food Competition task, where a small conflict for access to a discrete number of palatable pellets was introduced in the homecage of non-deprived adult male Sprague-Dawley cage mate rats. We decided to focus our study on this specific population as most of the available literature on dominance and aggressive behavior in rats has been performed in adult males ^13–17^ being Sprague Dawley rats among the most used laboratory strains in behavioral neuroscience. In order to validate this new tool, we compared the dyads’ behavior to that observed in other competition tests: 1) competition for 1% sucrose solution, 2) modified standard tests used in the field, that involve deprivation, where animals compete for food or water and 3) the Tube Test. We performed a detailed analysis of behavior in each test, and although no aggressive interactions were observed, our results indicate that stable hierarchies in rats are indeed detectable by the modified Food Competition test which is especially suited for their identification, based on its trial structure and the degree of conflict induced.

## Results

### Behavioral profiles differ across the social competition tests used

To identify social status within pairs of cage mates we performed the modified Food Competition test and other behavioral tasks in which the animals needed to compete for resources, either for palatable pellets, sucrose solution, water or tube occupation (Figure 1 and Supplemental Figure 1). All the tasks, except for the Tube Test, were performed in the animals’ home cage. Animals displayed different behavioral profiles depending on the configuration of the test, whether it had a trial structure, the amount of reward available, and their internal state (satiated vs deprived) (Figure 2 and Figure 3). In order to control for possible effects of winning history we created two independent groups where we counterbalanced the order of the tests. No differences were observed between the groups, suggesting that hierarchy was already established (Kruskal-Wallis test comparing the duration of consumption in the two counterbalanced groups: modified Food Competition test (mFC) Day1: X^2^(2)=0.026, p=0.871; mFC Day2: X^2^(2)=0.007, p=0.935; SC Day1: X^2^(2)=1.516, p=0.218; Sucrose Competition with continuous access to reward (SC) Day2: X^2^(2)=1.904, p=0.168; Sucrose Competititon with Intermittent access to reward (SCI) Day1: X^2^(2)=0.457, p=0.499; SCI Day2: X^2^(2)=2.055, p=0.152; mFCD: X^2^(2)=0.00, p=0.989; Water Competition (WC): X^2^(2)=0.293, p=0.589; Tube Test (TT) Day1: X^2^(2)=0.00, p=1.0; TT Day2: X^2^(2)=0.00, p=1.0). Data from both groups was thus merged for the rest of the analysis.

**Figure 1.**
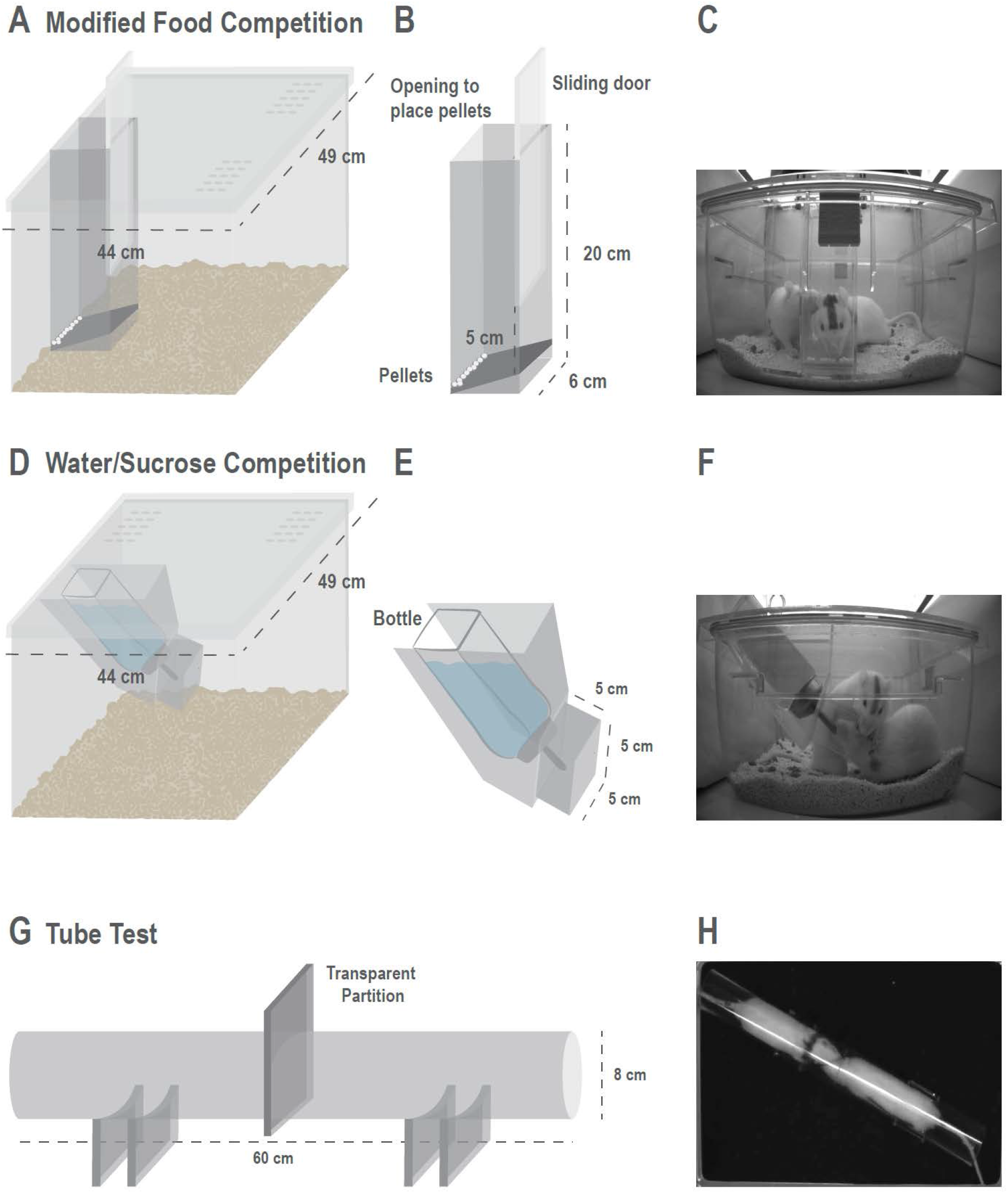
Design and measurements of the behavioral tests used for identification of established dominance status. **(A)** Schematic illustration of the transparent lid and feeder in the modified Food Competition apparatus. **(B)** Detailed schematic illustration of the feeder used in Food Competition protocols (with and without deprivation) including the measurements to fit the bottom part of a Rat IVC cage (Sealsafe PLUS Green Line ventilated cages, Techniplast). The sliding door can be opened leaving a 5 cm high access which only allows one animal to eat at a time, and allows a trial structure for the task. A small opening on the top of the feeder allows to refill new pellets during inter trial interval. If adaptation of measurements to another type of home cage is needed, we advise to leave 3-4 cm from the end of the feeder and the bottom of the home cage. This prevents bedding to go into the feeder, which difficults the visibility of the pellets while animals are consuming. **(C)** Two cage mates can be observed at the feeder area, where one is consuming the pellets while the other is pushing to get access to the food. **(D)** Schematic illustration of the lid and bottle holder for the Water and Sucrose Competitions protocols (with continuous or intermittent access) protocols **(E)** Detailed schematic illustration of the bottle holder. A 5×5 cm restraining tube around the lick spout was created to prevent simultaneous access of both animals to the resource. **(F)** Two rats behaving in the Water Competition task, where one of the rats is drinking while the other is pushing to have access to the bottle **(G)** Schematic of the Tube Test with measurements used in this task, the transparent partition in the middle of the tube is removed at the beginning of a trial once both animals reach this area. Laser-cut acrylic holders were used to give stability to the set up **(H)** Two rats interacting inside the tube during the initial moments of a trial.

**Figure 2.**
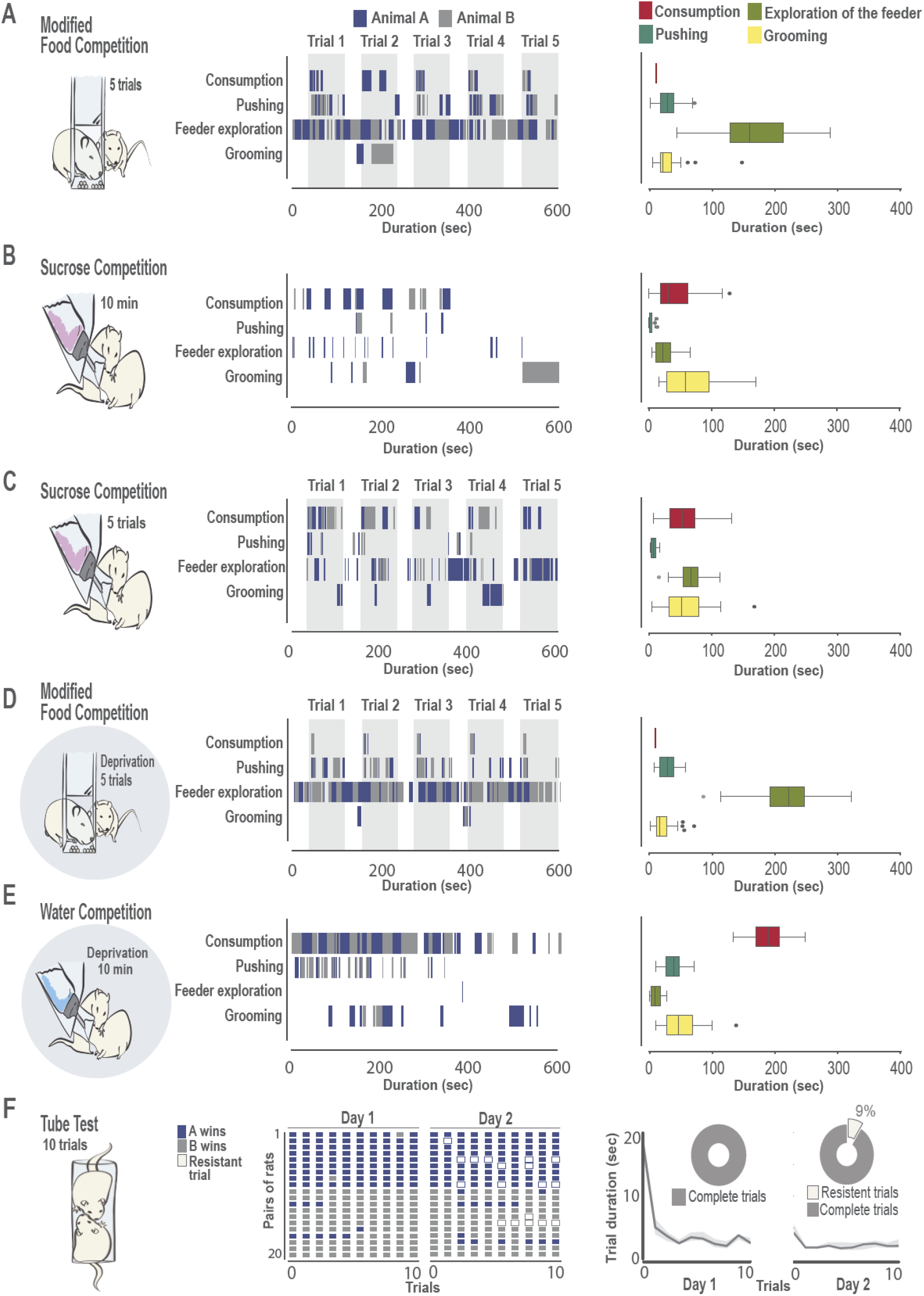
Behavioral profiles change according to the specifics of each social competition test. (**A**) modified Food Competition test, (**B**) Sucrose Competition with continuous access to the bottle, (**C**) Sucrose Competition with intermittent access to the solution, where animals could only drink during 2 minutes and the bottle was absent during 1 min inter-trial interval; (**D**) modified Food Competition test with deprived animals, (**E**) Water Competition test, (**F**): Tube test. For all tests, **left panel** shows the schematic representation of the task. Cartoons with a shaded circle as background indicate that tests were performed under deprivation. **A-F middle panel**: raster plots showing frequency and duration of behaviors of interest in an example pair of animals (animal A in blue, animal B in grey), except for **F** where the animal wining each trial is depicted for the 20 pairs of animals in all trials. White coloured trials in the Tube Test correspond to “resistant trials” where the loser of the pair resisted to enter the tube and trial could not be completed. In those tasks with a trial structure, (**A**, **C**, **D**) grey shaded areas indicate the moments where access to the resources where available, while intertrial interval, during which animals could explore the feeders but not access the food or sucrose bottle, are indicated with white background. **A-E right panel**: Boxplot representation of the duration of each behavior of interest for each individual animal. When more than one day of testing was performed, data represents average of the two days. For those tests with a trial structure, values represent behavior throughout the entire test period averaged over the 5 trials (or 10 trials, when two days of testing were performed), being one trial defined as the last 40 seconds of the intertrial interval and first 80 seconds of access to reward. Median, quartile 1 and 3 are represented, whiskers indicate minimum and maximum values and extreme values are signaled as a dot. Consumption in red, pushing in dark green, Exploration of the Feeder in light green, Grooming in yellow. **F right**: Trial duration in the Tube Tests in the two testing days. Black line represents the median duration that animals took to push their partner out of the tube, and grey shadows depicts 95% confidence interval. As an insert in each day, pie charts represents the percentage of trials that were completed. Note that 9% of trials in the second day of the Tube Tests were not completed as one animal (the loser) refused to enter the tube.

**Figure 3:**
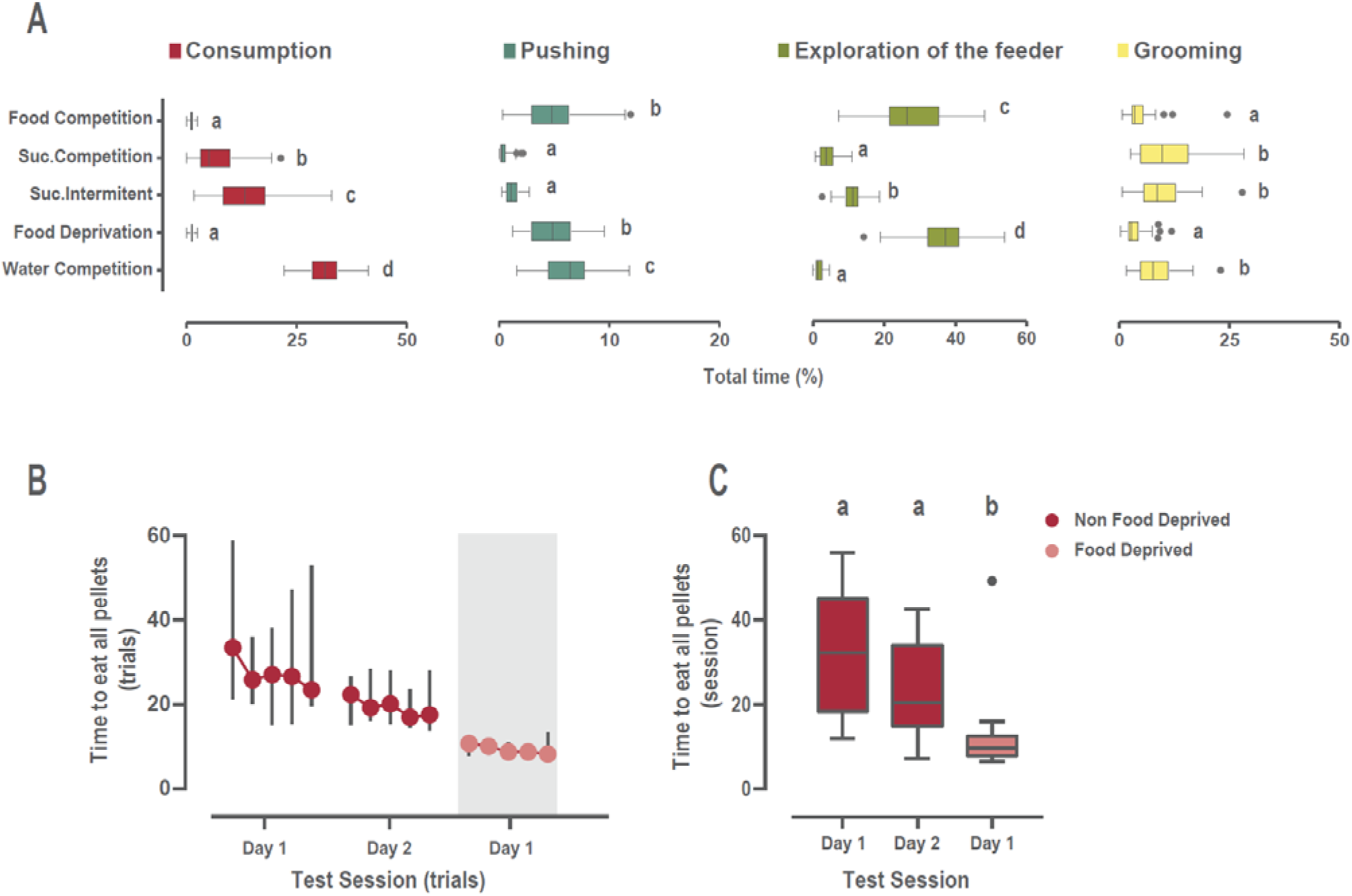
Descriptives of the behavioral analysis for all social tasks involving competition for resources. (**A**) Comparison of the behaviors of interest displayed by each animal across all behavioral tests. Data is presented as percentage of time performing a specific behavior related to the total duration of the task, enabling comparison between tests differing on duration. For those tasks with a trial structure, consumption corresponds to the first 80 seconds of reward availability for each trial, while pushing, exploration of the feeder and grooming durations correspond to the displays performed during the 40 seconds before and 80 seconds after reward availability. When more than one day of testing was performed, data represents average of the two days. Each competition test provided very different behavioral profiles, where the duration of consumption and exploration of the feeder or pushing behaviors to access the resources differed clearly over the tests. Note that pellets in the modified Food Competition test were consumed during the first seconds, but still high levels of pushing and exploration of the feeder where observed. However, the time spent consuming sucrose and the levels of pushing for accessing the bottle were low. Boxplots depict the median and quartile 1 and 3, whiskers indicate minimum and maximum values and extreme values are signaled as a dot. Consumption in red, pushing in dark green, Exploration of the feeder in light green, Grooming in yellow. Letters denote statistically significant differences between behavioral tests after one-way ANOVA with Tukey posthoc comparisons. Time spent grooming was not normally distributed, thus non-parametric analysis was performed. (**B**) Latency to eat all pellets for each trial is shown in the Food Competition tests, with or without deprivation where median and 95% CI for each trial are represented. Animals only took around 20 seconds to eat all pellets available, decreasing this latency when food deprived. Shaded background indicates session performed under food-deprivation. (**C**) Boxplots representing the average time to eat all pellets over testing days. Wilcoxon rank test revealed that latency to eat the pellets decreased on the second day of testing of the modified Food Competition test, and did further under deprivation state. **p<0.01, ***p<0.001

In the modified Food Competition task (Figure 2A, Figure 3, **Movie 1**), the limited number of available food pellets (10 per trial) led to a very fast consumption of resources which lasted a few seconds (Figure 3 B-C). Interestingly, although pellets were consumed in the first seconds of each trial, animals displayed high levels of exploration of the feeder and displayed notable amounts of pushing, suggesting high expectation of reward (Figure 3A). On the contrary, during the Sucrose Competition task, exploration of the reward location and pushing levels were low (Figure 2B and Figure 3A), but exploration of the bottle location increased when access to the sucrose bottle was presented in an intermittent manner (Figure 2C and Figure 3A).

As expected, modulation of internal state (food or water deprivation) affected the behavior of the animals. In the modified Food Competition task with deprivation (Figure 2D) animals consumed the pellets faster (Figure 3C, Wilcoxon signed rank tests against non-deprived Food Competition day 1: z=-3.659 p=0.0003; day 2: z=-3.136 p=0.002) and spent significantly more time investigating the feeder (Figure 3A), although the amount of pushing did not differ from that displayed in non-deprived animals (Figure 3A). When competing for access to water under deprivation (Figure 2E), animals dramatically increased the time they spent drinking compared to consumption displayed in the other tests (Figure 3A), performing long bouts of drinking and alternating between animals (Figure 2E). Levels of exploration of the water bottle were low compared to the rest of the tests. Surprisingly, although the motivation to drink was high, revealed by the long water consumption time, pushing levels did not increase proportionally (Figure 3A).

Finally, the dyads established a very stable winner/loser relationship in the Tube Test (Figure 2F), where most of the animals that would start winning in the first trials would continue doing so over the remaining trials. This winner/loser structure was maintained across the two testing days (Figure 2F **middle panel,** raster plot of winning history for all pairs). Interestingly, on the second day of testing, the loser partner of some pairs showed reluctance to enter in the tube (9% of the total trials), suggesting a strong subordination towards the partner. The time to solve the conflict in the Tube test, measured as the latency from the moment the partition at the center of the tube was removed until one of the rats was pushed out of the tube, rapidly decreased after the first trial, reaching then fast and stable latencies of around 3.5 seconds on average (Figure 2F, right panel).

### Social hierarchy as priority access to resources

We categorized the animals of each pair as dominant (D) or submissive (S) according to the amount of resources they would consume within each test (pellets eaten in the Food Competition tests, time spent drinking in the Sucrose or Water Competition tests, and the number of wins in the case of the Tube Test). According to this criterion, as expected, animals categorized as dominant consumed significantly more resources than their partners in every task (Figure 4A-E) (Wilcoxon signed-ranks for consumption in mFC: z=-3.92, p<0.0001; SC: z=-3.82, p<0.0001; SCI: z=-3.92, p<0.0001; mFCD: z=-3.83, p<0.0001; WC: z=-3.92, p<0.0001). In the same line, one animal always won more encounters than the other in the Tube Test (Figure 4F, Wilcoxon signed-ranks for TT Day1: z=-4.06, p<0.0001 and for TT Day2: z=-4.12, p<0.0001), with the exception of one pair of animals in Day 1 and another in Day 2, where both animals of the pair won the same number of trials, thus no categorization as dominant or submissive was possible in these cases.

**Figure 4.**
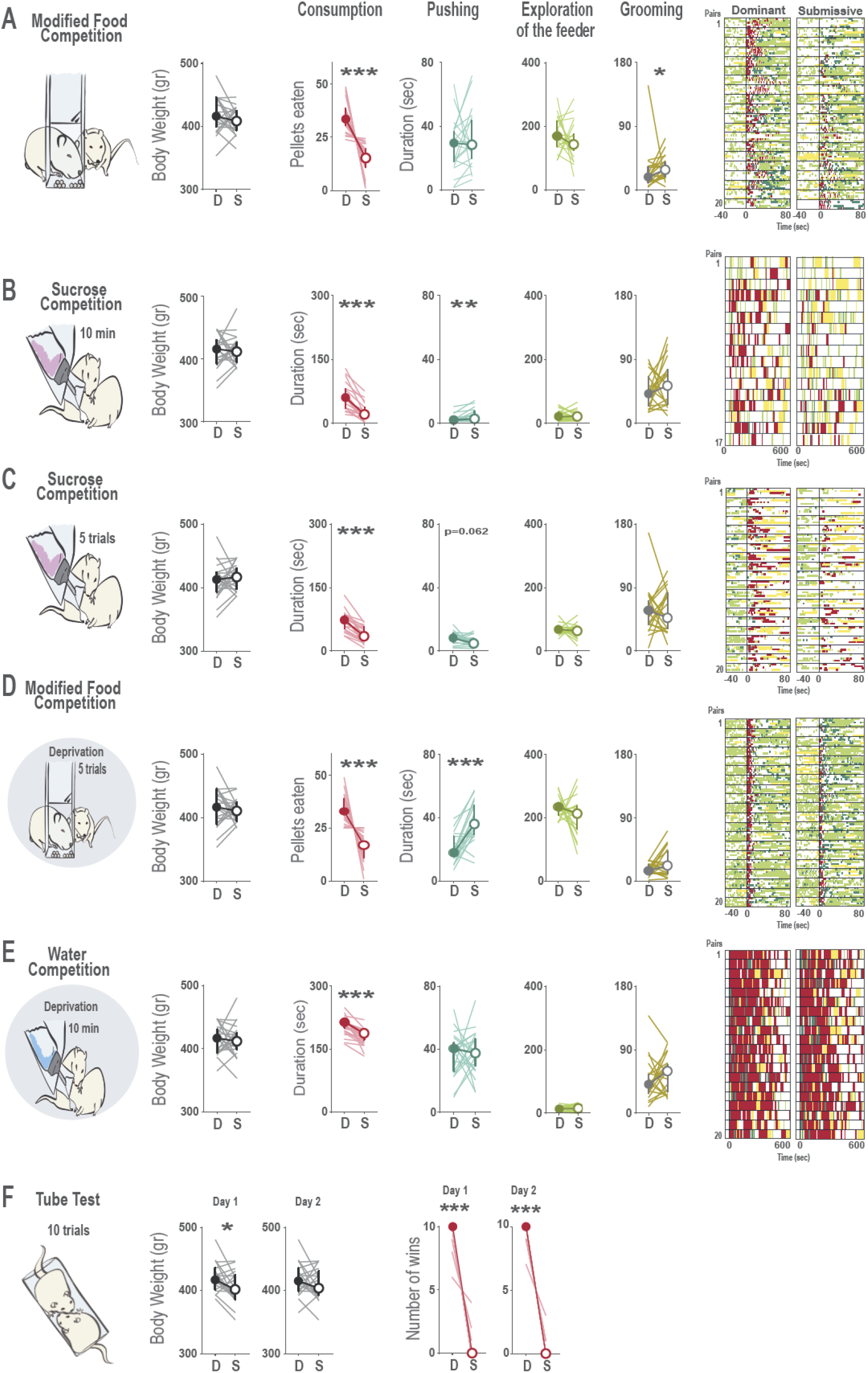
Categorization of animals as dominant and submissive according to behavior within each task. (**A-E)** Behavioral profiles when consumption of the resources within each test is used to define dominant and submissive animals. For all panels, a schematic cartoon with experimental design is provided. Cartoons with shaded background indicate that tests were performed under deprivation. For each test, differences between dominant and submissive animals are represented regarding body weight, time spent consuming the resource (or number of pellets eaten in the case of the food competition tasks), duration of pushing, exploration of the feeder and grooming is provided, where median and 95% interval confidence are displayed and individual values are showed with light lines. Color coded raster plots of behaviors of interests display raw data in a testing session, where pairs of animals are sorted according to the stronger differences in hierarchy for each test. For those tests with a trial structure, the five trials of each pair are plotted in separate lines, all aligned to time 0 (when access to reward was possible) and reflecting the behavioral data of the 40 seconds before and 80 seconds after that moment. When more than one day of testing was performed, data represents average of the two days and rasters display data of day 2. In those cases where the animals of one pair had identical values in the categorizing criteria (same amount of pellets eaten, duration of consumption or trials won in the Tube Test), no hierarchy was assumed for this specific task, and these pairs were removed from this analysis (one pair for the Sucrose Test, one for the Tube Test and one for the Food Competition under deprivation). For all graphs and rasters, consumption is represented in red, pushing in dark green, exploration of the feeder in light green and Grooming in yellow. Time of consumption in each test was significantly different between animals defined as dominant or submissive, and in some cases, these differences also were translated to a differential amount of grooming or pushing behavior. Interestingly, no differences in body weight were found in these tests, indicating that priority access to resources in established hierarchies is not influenced by the size of the animals. (**F)** In the Tube Test, the amount of winnings was clearly different between dominant and submissive rats, and was related to differences in body weight, which reached significance in the first days of testing, being bigger animals those more likely to win. *p<0.05, *\*\*p<0.01, *** p<0.001*.

We decided to investigate how body weight would relate to dominance in established hierarchies of rats. Intriguingly, we did not observe differences in the body weight between dominant and submissive animals in any of the behavioral tests where animals would compete for food, sucrose or water (Figure 4 A-F; Paired Sample T-test for weights in mFC: t=1.208, p=0.242; SC: t=-0.309, p=0.761; SCI: t=-0.843, p=0.410; mFCD: t=0.522, p=0.608; WC: t=0.067, p=0.947). In contrast when they had to compete for territory for the first time in the Tube Test, a significant relationship between dominance and body weight was observed (Figure 4F; Paired Sample T-test for weights in TTDay1: t=2.529, p=0.021). In conclusion, although no relation was observed between social hierarchy and body weight in the rest of the tasks, this was not the case in the Tube Test, where bigger rats had a higher probability of winning in the first encounters.

We then asked whether hierarchy following this criterion, amount of resources consumed, would also translate to differences in other behaviours within each test. General exploration of the resource location during the whole session did not differ between dominant or submissive animals when consumption in the same test was taken as criterion. However, we did observe that submissive animals would spend more time self-grooming in the modified Food Competition test (z=-2.016, p=0.044) and that time spent pushing was modulated by dominance in some tests. Dominant animals tended to display more pushing in the Sucrose Competition with intermittent access (z=-1.867, p=0.062) and surprisingly, submissive animals were the ones that pushed more in the Sucrose Competition with continued access to the bottle and the Food Competition under deprivation (SC: z=-2.722, p=0.006 and FD z=-3.472, p=0.0005). As this observation was unexpected, we next explored this further.

### Dominant rats are more efficient displacing their partners to gain access to resources

Although the time animals spent pushing their partner should be a good measure of amount of conflict between the interacting animals, qualitative differences might be more informative of dominance status. One possibility is that even if a dominant rat pushes less often, its thrusts may be more successful in removing the partner from the resource. Thus, we calculated for each animal the percentage of successful pushing from the total number of pushing bouts, i.e. the fraction of pushing epochs that actually displaced the partner and allowed access to the resource.

Strikingly, dominant animals were more successful in displacing their partners in the modified Food Competition under deprivation, while submissive animals would push often but failed to displace their partner (Figure 5, Wilcoxon signed rank test z=-1.979 p=0.048). However, this was not observed in the Water Competition test (WC: z=-0.821 p=0.411) nor in the tasks not involving deprivation (mFCD: z=-0.933 p=0.351; SC: z=-1.014 p=0.310; SCI: z=-0.563 p=0.573). The lack of differences in successful pushing between dominant and subordinate rats in the water competition test, could result from a limited window within the test where asymmetric interactions are apparent. To investigate this possibility, we identified the epochs with highest conflict in the Water Competition test, i.e., those where the most drinking and pushing behavior was observed for each pair of animals (Supplemental Figure 2A). We then classified the animals as dominant and submissive according to the duration of drinking in that epoch and quantified pushing displayed by either dominant or subordinates. This new categorization led to a change in the hierarchy in 55% of the pairs (11 out of 20). Here, although submissive animals spent longer time pushing their cage mate (Figure 5E middle plot, z=-3.920, p<0.0001), again dominant animals displayed higher efficiency in displacing their subordinates, being successful practically 100% of the times (Figure 5E plot on the right, z=-3.627 p=0.0002). This fine grained behavioral analysis, where pushing is categorized into successful or unsuccessful, thus revealed that although in some tasks submissive animals displayed higher duration of pushing, they rarely managed to get the access to the resource, being dominant rats more successful to displace their cage mates.

**Figure 5.**
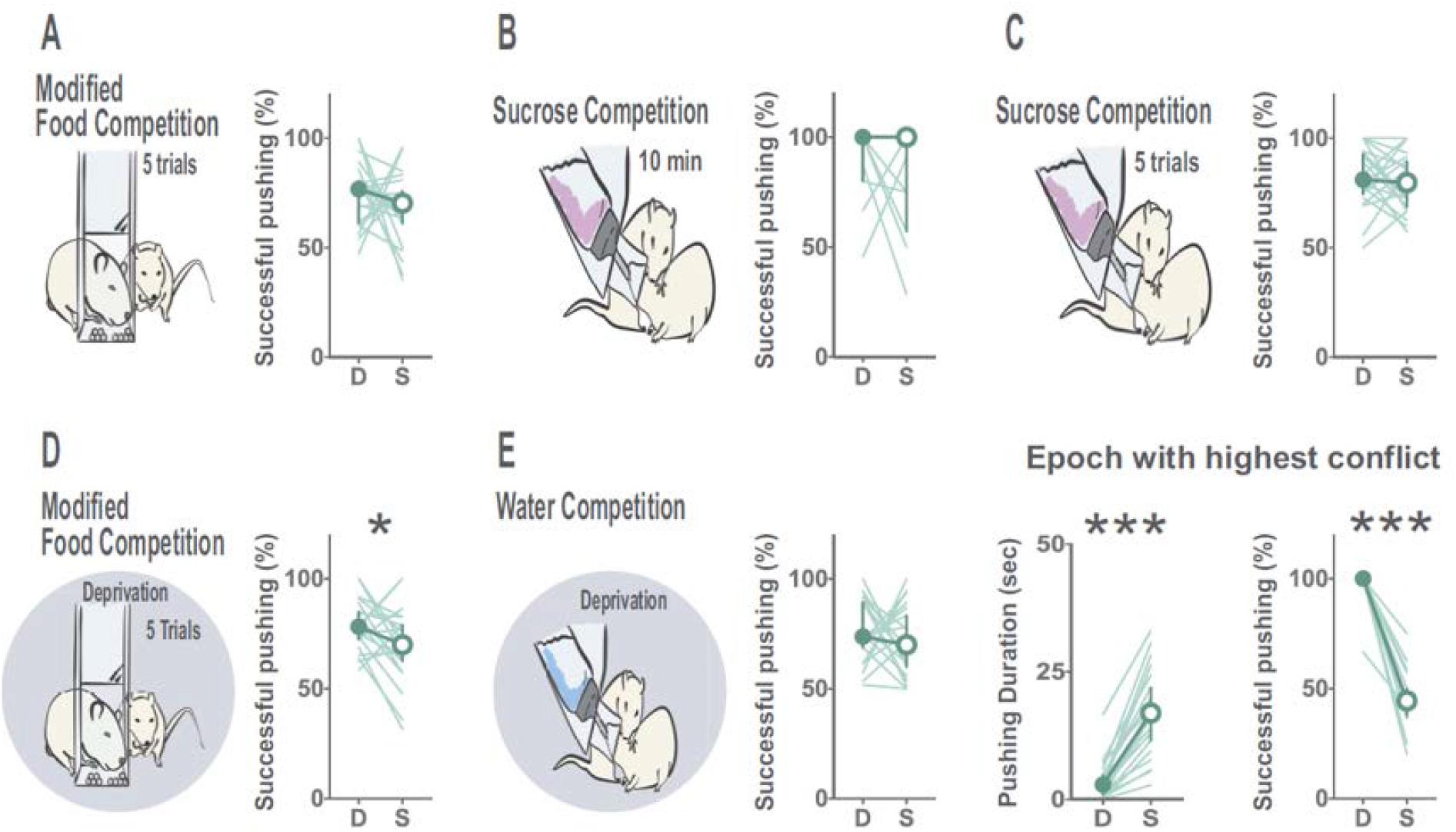
Dominant animals are more successful in displacing their subordinate partners to gain access to resources when in deprivation states. Dominant animals showed a higher percentage of successfully pushing away their subordinates while gaining access to the resources in moments with high conflict. Percentage of successful pushing did not reach significance in the modified Food Competition (**A**), Sucrose competition with continuous (**B**) or Intermittent access to the bottle (**C**) but was significantly different in the Food Competition under deprivation (**D**). No differences were observed in the efficiency of pushing behavior in the Water Competition task (**E**). However, when assessing these differences in the moments with highest conflict in this test, defined as the bout where intense drinking was displayed and high levels of pushing behavior were observed, submissive animals spent more time pushing, but dominant animals were almost always successful to displace their partners in every bout of pushing. D: dominant, S: submissive according to the consumption in each test. Median, 95% CI and individual values for all animals are represented. *p<0.05, *** p<0.001.

### Social hierarchy as a stable trait between tests

To examine whether social hierarchy is a stable trait in familiar animals, we analyzed reliability across the performed tests. To this end, we computed the *Dominance Index* (DI) for each test where the difference in resource consumption across partner animals was normalized by the summed resource consumption of the pair. A DI close to 0 means that animals did not have a strong hierarchy. Positive values indicate that animal A consumed more, while negative values indicate that animal B was the one having priority access to resources or won more trials in the Tube Test. Provided that behavioral measures were stable across testing days (Supplemental Figure 3) data was averaged for this analysis.

DI for positive reinforcers led to a highly variable distribution across dyads, where in some dyads animals would strongly differ in their consumption and in others differences were subtle (Figure 6A). This was not the case in the Water Competition, where DI was mostly around 0 for all animals, indicating that both animals drank very similar amounts of water during the test. In contrast, the Tube Test gave very polarized DIs, where most of the pairs had one animal winning almost 100% of the trials. To refine our understanding of the hierarchy in the Tube Test, we computed a *Conflict Resolution Index (CRI)*, which would take into account not only who won a trial, but also how long it took for the conflict to be resolved (i.e., the latency for one of the animals to be pushed out of the tube). The conflict resolution index revealed a more continuous and fine-grained measure of hierarchy strength in each pair (Figure 6B).

**Figure 6.**
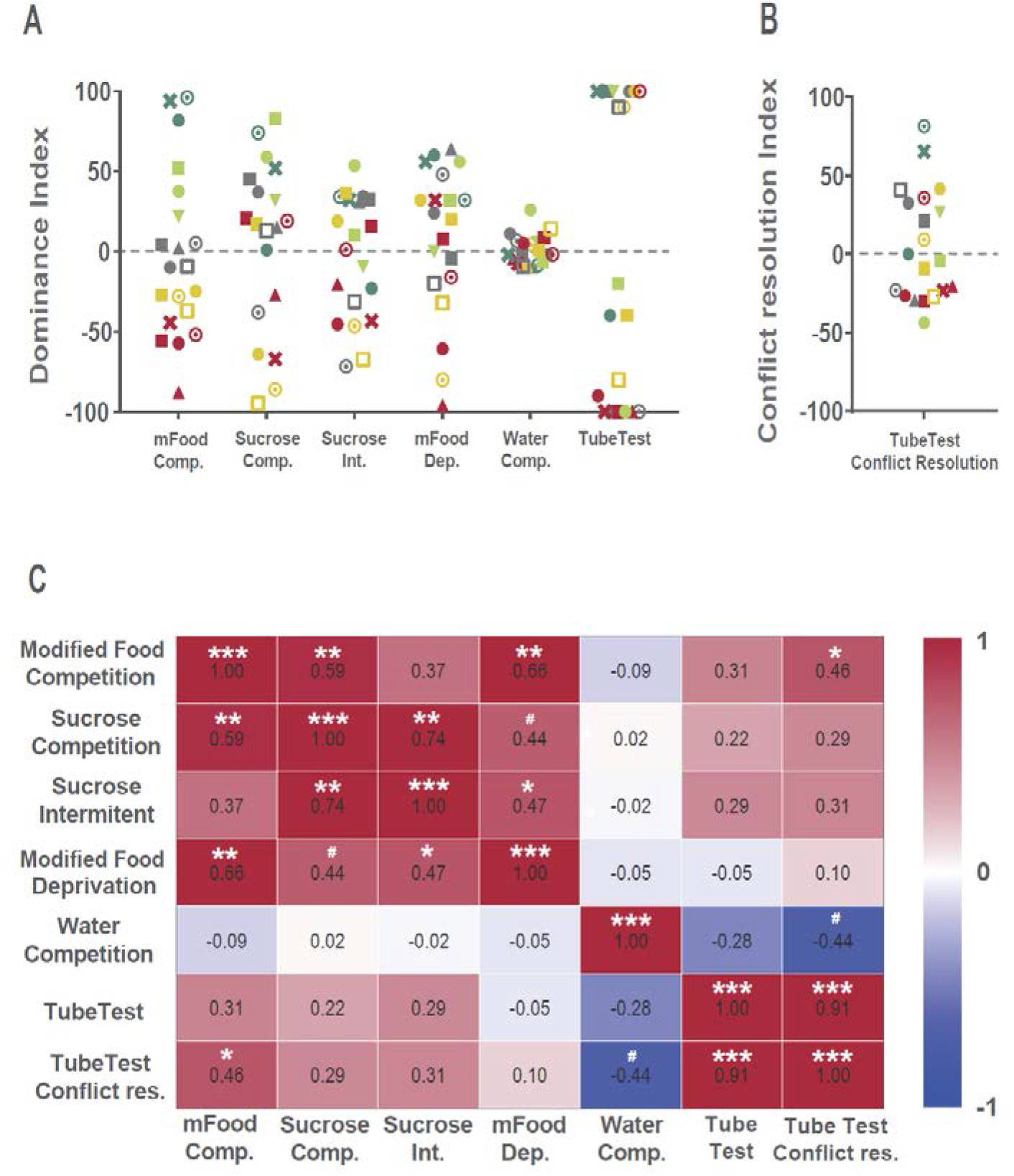
Reliability between dominance measurements between tests. (**A)** Dominance index (DI) based in consumption of resources for each behavioral test is represented being each pair of animals identified with a specific color/symbol. In those cases where two days of testing was performed, values plotted correspond to the average DI of the two days. Competition for positive reinforcers produced highly variable DI indicating detectable differences in the strength of social hierarchy between the pairs. Water Competition, however produced DI close to 0 indicating that the animals of most of the dyads drank very similar amounts during this test. The Tube Test produced a very polarized distribution of DI, where one animal of each pair would win most of the trials. (**B**) These differences in hierarchy became less polarized when taking into account the time the animals took to solve the conflict in the tube (DI in the Tube Test /latency to finish the trial). (**C)** Correlation matrix between DI from all tests. Competition for positive reinforcers DIs were correlated across tests, and DI from the modified Food Competition test correlated with the Tube Test, when conflict resolution time was taken into account. However, no significant correlations were observed with the Water Competition tests. #p<0.10, *p<0.05, **p<0.01, *** p<0.001.

If social hierarchy was a stable trait, then dominance indexes should be correlated across tests. We found DIs from tests with competition for positive reinforcers were positively correlated (Figure 6C). Moreover, the modified Food Competition test correlated with the DI index in the Tube Test, when conflict resolution time was taken into account. Water competition DI was not correlated with any of the other tests. Water deprivation could have challenged the homeostasis of the interacting animals bringing them to a very different internal state that disrupted the stable hierarchies revealed by the other tests. Alternatively, computing the DI for the water competition test using the whole test duration, may have diluted differences in water consumption across animals within the pair. Hence we re-calculated DI for different time windows, just as we observed for the pushing behavior (see above). No correlation was found between water competition test and the other test when other time windows were used to compute DI (Supplemental Figure 2B). Still, we have shown that during this test, the dominant animals accessed the water bottle in a qualitatively different manner, by successfully displacing their subordinates (Figure 5E). This highlights the importance of considering multiple behaviors simultaneously and suggests that social status in the Water Competition task could be better assessed by finer behaviors rather than water consumption.

### Modified Food Competition tests as a tool to measure stable hierarchies

Last, we calculated for each task a *Conflict Index* as a measure of the degree of conflict that our manipulations introduced in the home-cage, by dividing the time animals spent pushing in one test by either the time spent consuming in case of the Sucrose and Water Competitions, or latency to eat all rewards in the case of the modified Food Competitions. Food competition with and without deprivation were the tasks with higher conflict, as animals displayed high amounts of pushing and the time available to eat the resources in each trial was very short (Supplemental Figure 4A). Since the modified Food Competition test yielded significant levels of conflict but did not involve food deprivation, we next evaluated whether attributing dominance within dyads using this test, would allow correct identification of the dominant rat in the other tests. Specifically, animals were classified as Dominant or Submissive according to their DI in the second day of testing in this task, as the conflict index was higher in this day (Supplemental Figure 4B). Those pairs where the difference of the number of pellets eaten was small (less than 5% difference compared to equal consumption between the animals) were considered to have an unstable or unclear hierarchy (n=4) and were not included in this last analysis. As expected, dominant animals consumed more pellets in the Food Competition test (Figure 7A, Wilcoxon signed-ranks of average Consumption in mFC of both days z=-3.362, p=0.001). Interestingly, they also successfully pushed their partner away from the feeder more (z=-2.275, p=0.023), explored the feeder more during inter the trial interval, where the pellets were present but not accessible (z=-2.844, p=0.004), and groomed less than their submissive pairs during the trial period (z=-2.275, p=0.023). Moreover, dominant animals according to the modified Food Competition did consumed more sucrose, both when access was continuous (Figure 7B, SC z=-2.499, p=0.012) or intermittent (Figure 7C, SCI z=-2.275, p=0.023). Attributing dominance found in the modified Food Competition, to the same test run under food deprivation, revealed similar dominance interactions (Figure 7D, Consumption in mFCD z=-2.619, p=0.009; Successful Pushing in mFCD z=-2.534, p=0.011; Anticipatory exploration of the feeder in mFCD z=-2.902, p=0.034, Anticipatory grooming in mFCD z=-1.992, p=0.046). Moreover, dominant animals also successfully pushed away their partner more in the Water Competition test (Figure 7E, z=-2.379, p=0.017). In the case of the Tube Test, the amount of wins did not differ between dominant and submissive animals, as determined by the modified food competition test (Supplemental Figure 4C). In figure 3 we show that differences in body weight affects the probability of winning in the Tube Test, especially on the first day. This was however not the case for the other tests. Thus, weight may dominate the outcome of the tube test, overshadowing the dynamics of social interactions within stable pairs. Therefore, we decided to examine the relationship between the hierarchy in the modified Food Competition and the Tube Test while controlling for the effect of the body weight. To this end, we first regressed the number of pellets eaten against the animals’ body weight and calculated the residuals. Next, we calculated, in the same manner, the residuals when regressing out body weight from the conflict resolution index (see above) of the first day of the Tube Test. Interestingly, the linear regression of these residuals was statistically significant (p=0.010) indicating that indeed, when correcting for the effect of body weight, consumption in the modified Food Competition predicts who will win in the first interactions of the Tube Test (Figure 7F). Thus, dominant animal in the modified Food Competition test also won more trials in the Tube Test when the influence of the body weight was controlled for.

**Figure 7:**
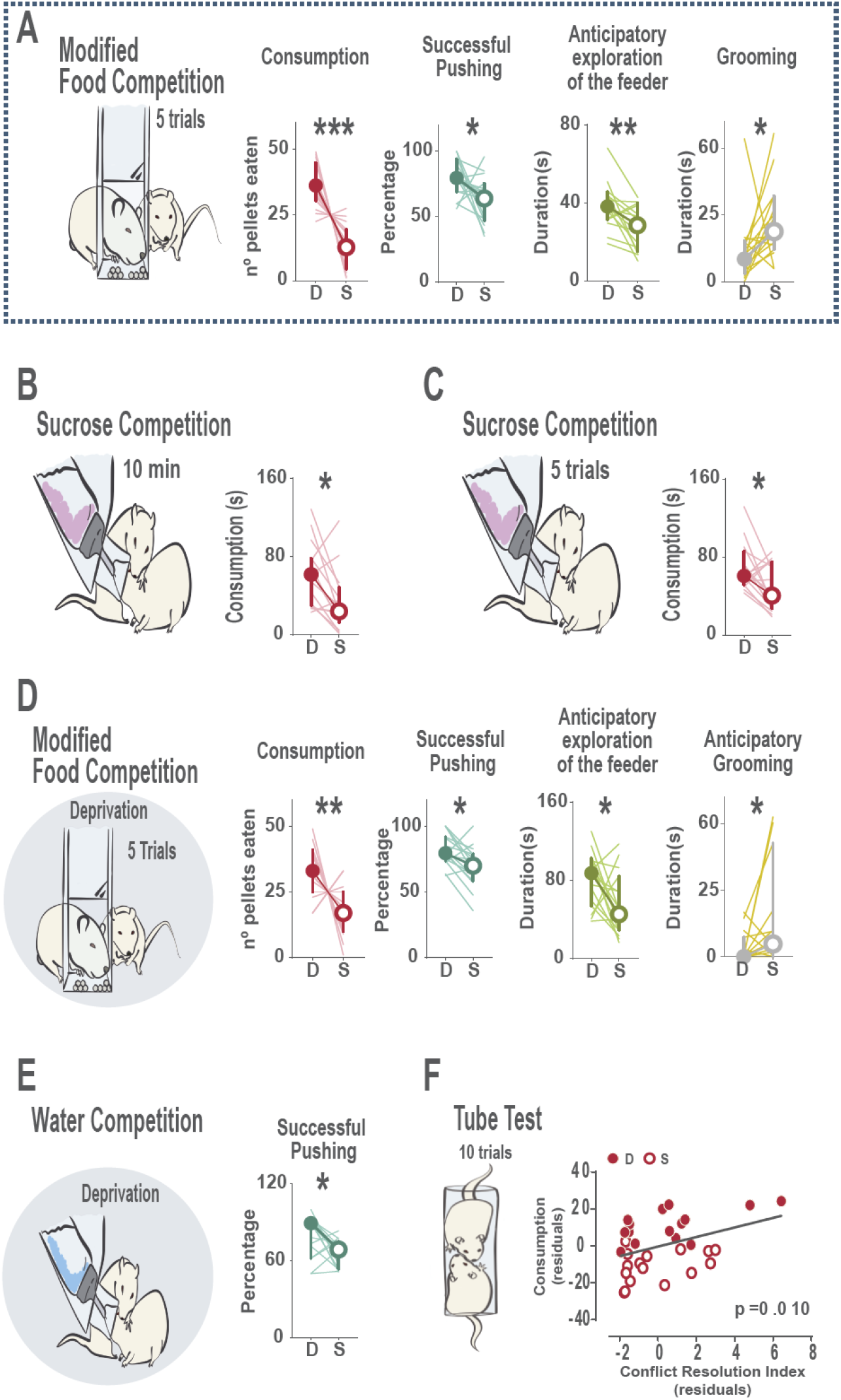
Food competition in the home cage is a simple and reliable measure of established social hierarchies in rats. Animals displaying differences in the amount of pellets eaten in the modified Food Competition test were classified as dominant or submissive animals, and their behavioral profiles studied in the rest of the tests. (**A**) When considering average behaviors of both testing days in the modified Food Competition test, dominant animals significantly ate more pellets, were more successful to displace their partners from the food magasin during the competition and explored more the location of the resource when access was still prevented, and groomed less time throughout the session (considering the last 40 seconds of intertrial interval and first 80 seconds of reward exposure of each trial). Moreover, they (**B**) drank more in the Sucrose Competition test with continuous, and (**C**) with intermittent access to the bottle. (**D**) Similar behavioral profiles were displayed in the Food Competition under food-deprivation, where differences in consumption, successful pushing, anticipatory exploration of the feeder and grooming were also observed. We defined the anticipatory window as the last 40 seconds of the intertrial interval, just before the bottle was placed in the lid thus consumption was still not possible (**E**) Dominant animals according to the modified Food Competition test, were also more successful to displace their submissive in the Water Competition test, indicating that although no differences in the amount of water drank were observed, the quality of the interaction was significantly different. (**F**) The amount of pellets eaten in the modified Food Competition test significantly predicted the probability of wining in the first interactions in the Tube Test, when latency to win was taken into account, and values were regressed out of the influence of body weight. Dashed line around panel A indicates that consumption in second day of testing in the modified Food Competition test was the criteria to evaluate differences in all the tasks. Median, 95% CI and individual values for all animals are represented. * p<0.05, **p<0.01, ***p<0.001.

## Discussion

Here, we developed and validated the modified Food Competition task, a new tool designed to provide, for the first time, the means to evaluate established hierarchies in pairs of rats, and with the added value of doing so in the home cage, without inducing aggressive behaviors nor requiring food-deprivation.

In our task, cage mates with a stable hierarchy competed for access to positive reinforcers. The introduction of a small conflict in the home cage, where consumption of appetitive food is only possible for one animal at a time, led to subtle competition which translated into increased consumption by the dominant rat. This measure reliably predicted differences in other behaviors observed during the competition test. These differences held across the different social tasks evaluated, such as competition for sucrose solution or food and water competition under deprivation. Importantly, testing was performed in the home cage of the animals, thus minimizing the influence of the experimenter in the social interactions displayed, and interference by other factors such as anxiety or exploratory behaviors usually displayed in novel environments. Conflict in the modified Food Competition increased over days, providing the second day a clearer picture of the established hierarchy, most probably because during the first of competition in the home cage, differences in attention to the appearance of the conflict situation might be modulating the interactions. In future experiments, it might be desirable to include a third day of testing to confirm whether conflict continues increasing or is stable after the second day.

Interestingly, although competition was observed and hierarchy could be identified, no agonistic behaviors (biting, boxing, keep down, lateral threat) were observed between the cage mates in our task. This is in accordance with previous reports indicating that once social hierarchies are established, the number of agonistic encounters decreases ^6^. Moreover, the fact that no food deprivation was required, nor aggressive behaviors were observed, can be considered as an added value of our task. Aggressive behaviors induce robust stress reactions in rodents ^20^, and food deprivation, although widely used in neuroscience to increase motivational salience during behavioral testing, modifies internal state ^21^ and social behavior ^22^. Minimizing the possible long term effects of these manipulations on the internal state of tested animals is particularly important for studies regarding the impact of social hierarchy on other behavioral, physiological and brain functions.

Moreover, the fine behavioral characterization across the different social tasks used, allowed us to identify very interesting social patterns, that to our knowledge have not been reported before. In this direction, we describe that social hierarchy tasks with a trial structure (modified Food Competition tasks and Sucrose Competition with Intermittent access), where access to resources was presented in a repeated and intermittent manner, promoted competition. Moreover, we describe that when measuring conflict, the time spent pushing by the animals is not indicative of dominance, but rather how efficient an animal is to displace its partner while pushing. Although the dominance index in tasks that involved competition for positive reinforcers reliably revealed the strength of social hierarchy within a pair, this was not the case in the Water Competition test under deprivation, where the animals drank around the same amount of water (dominance index around 0), nor in the case of the Tube Test, where very polarized results were observed. In the Water Competition task, animals tended to perform long bouts of drinking, alternating consumption until both animals were satiated, which resulted in very similar final consumption levels in both animals. We asked whether an analysis with finer temporal resolution could unveil structured dynamics of water consumption in this task, such that asymmetries across rats in the dyad would emerge during bouts of conflict, and these asymmetries would correlate with other social competition tests. However, to the extent that we could quantify, we did not observe such a pattern. Strikingly, we found that although consumption between animals was very similar, other behaviors displayed while approaching the water bottle were clearly different. Although submissive animals in this task spent more time pushing their cage mate in moments of high conflict, they were rarely successful in accessing the water if the dominant was already drinking. Indeed, the ability of dominant rats to successfully displace their partners from the resource location was not limited to the Water Competition, being reliable in those tests where more conflict (pushing) was observed.

In the case of the tube test, we showed that computing conflict resolution index which takes into account multidimensional behavioral measures, such as the winner of a trial and conflict duration (latency for one of the animals to be pushed out of the tube) revealed a more granular view of the strength of social hierarchy in this test. In addition, this conflict resolution index correlated to the social hierarchy observed in the modified Food competition test, especially when taking into account the body weight of the animals. It is important to note that the final output of the tube test (i.e. who was the winner) did not correlate with other measures of social hierarchy. Although this test is widely used in mice (see ^3^ for review), our data indicates that in rats this test might not be an appropriate tool to measure established hierarchies, as who manages to push out of the tube its partner is largely affected by the body weight of the interacting animals. However, the conflict displayed inside the tube in the first encounters (incorporating the latency to win in the conflict resolution index), might be a better measurement of established hierarchies for rats. These results thus underscore the necessity of including multidimensional analysis of behavior and the importance of taking into account qualitative measurements when describing social interactions. Furthermore, although it might be surprising that the tube test is not a reliable measure of established hierarchy in rats, important differences in social behavior between mice and rats are starting to be reported ^22, 23^. Rats are more socially tolerant, and less hierarchical compared to mice. This might be related to their natural behavior in the wild, where rats are often observed in larger groups ^12^. These ecological differences should be carefully taken into account when borrowing tools from one species to the other ^24^.

Finally, body weight has been largely assumed to be a good indicator of social hierarchy in rats ^25^, although this has not been replicated in mice ^19, 26^. This view was inspired by classic works ^27^ and the seminal contributions in the field upon the development of the visible burrow system ^13^. However, our data, obtained in animals maintained with *ad libitum* access to food and water, did not support this observation, as dominant and submissive animals showed no differences in body weight, with the notable exception of the Tube Test. It is important to take into account that in our experiments the difference in body weight between the animals of a pair was lower than 10%, as they were age-matched. It is possible that larger body mass differences would indeed influence social rank in these tests, as previously shown in mice ^28, 29^ and rats ^25^. However, when using animals with marked body weight asymmetries, differences in age, and thus social experience, should be then taken into consideration as possible modulators of hierarchy. In our conditions, only the Tube Test was affected by body weight, where bigger animals had indeed more chances to win in the first encounters in the tube, probably indicating that in rats, as opposed to mice ^8^, body mass does affect the pushing behavior and the output in this task.

In summary, here we provide and validate a novel trial-based dominancy assay to be performed in the home cage of familiar non-deprived rats, the modified Food Competition test, which is easy to adapt and implement in any behavioral laboratory. The ability to assess dominancy in stable social hierarchies opens the possibility to study how social status affects different aspects of an individual cognitive, behavioral and physiological functions in the context of various social interactions, regarding which very little is known. Intense efforts in the last years have highlighted the Norway rat as a very interesting animal model to identify the proximate mechanisms and neural circuits of complex social functions ^30–42^. It is conceivable that each of these behaviors is modulated in some way by hierarchy.

While most of the available tests for the evaluation of social hierarchy in rats are based on the identification of dominant animals during the establishment of a new hierarchy ^3, 11, 15, 16, 26^ it is uncertain whether becoming the dominant in a first encounter will translate into keeping the same rank when the hierarchy becomes stable. Indeed, previous reports indicate that repeated encounters are needed for two unfamiliar animals to establish a stable social rank ^14–16^. By studying social status only during the early phases of its development, we are losing a huge aspect on the richness of social behavior and how it might be impacting brain function, in health and disease. The differences between the establishment of a hierarchy and its maintenance are largely unexplored, and the field would vastly benefit from new behavioral tools to address this fascinating question. Our new behavioral task opens the possibility for the study of such differences in rats.

Although we acknowledge that the general tendency in the field is to favor the use of mice as a model species, due to the large advantages related to its genetic tool box, cross-species validation is of utmost necessity, and validation of tools in different species an urgent need in Neuroscience. On the other hand, rats display much more sophisticated social and non-social behaviors compared to mice, and the development of tools such as CRISPR/Cas9 and state-of-the-art viral approaches, are making more accessible the precise monitoring and manipulation of neural circuits in other species. Taking all together, we foresee a drift towards a diversification of the species used in Neuroscience in the following years, and validation of ethologically relevant behavioral tools within the ecology of each species is needed.

Our modified Food Competition task provides a simple, robust, and unintrusive means of assessing established social hierarchy that can be readily incorporated into future studies, with the notable advantage of not inducing aggressive behaviors between the interacting individuals, nor having to manipulate internal state (deprivation), and being performed in the home-cage. One limitation of our work was the use of a very specific population for our study: male adult Sprague Dawley rats. Although we do not anticipate major problems in using the modified Food Competition test in other rat strains, behavioral differences have been already reported in several social and nonsocial behaviors between Long-Evans, Wistar and Sprague-Dawley rats ^23^. It could be possible that different rat strains would react differently to the subtle conflict we induce in the cage during the modified Food Competition test, and that acute aggressive encounters could be observed depending on the strain. Future studies could also expand our task to other developmental ages (such as peripuberty or late adolescence), where play-fight behavior and aggressive profiles are being acquired ^20^. Importantly, future studies should investigate whether social hierarchy could be assessed in female rats using the modified Food Competition Task. The neural mechanisms of female dominance and aggression have been surprisingly overlooked, and most of the knowledge in this direction has been obtained in the context of maternal aggression ^43–45^, with some notable exceptions ^46, 47^, moment where females display robust, clear and strong aggressive episodes. However, established social rank, contrary to *de novo* establishment of social hierarchies, is not based in aggressive encounters. We therefore believe our task is well suited for the study of social hierarchy in female rats, as dominance is evaluated not according to the quantification of agonistic behaviors, which female rats do not typically or strongly display, but on the behavioral response towards a subtle conflict to gain access to appetitive reinforcers. Moreover, although not the aim of the present work, our task can be easily scalable to larger groups of animals living in standard home-cages or more naturalistic environments. A collective evaluation of established hierarchies by introducing subtle conflicts during discrete periods in the home cage might provide very interesting information of complex social structures, and would suppose a clear advantage when evaluating groups of animals, as compared to repeated testing across multiple pairs, as the round-robin design currently used in the tube test in mice.

In conclusion, here we present a tool that allows, for the first time, to identify established social hierarchies in rats. After the precise description and validation of the modified Food Competition test presented here, this task is very easy and cheap to implement in any behavioral laboratory, which we expect will substantially help accelerate discovery on the effects of established social hierarchies on brain function. Our work adds to recent efforts towards the development of ethologically relevant paradigms ^48^, where rich information of the social interactions of rodents can be obtained with minimal intervention of the experimenters but in controlled and highly quantifiable laboratory settings.

## Materials and Methods

### Animals

40 three-months old male Sprague-Dawley rats (OFA, Charles-River, France) weighing 325-410 g at the beginning of the experiment were used. Upon arrival from the commercial vendor (Charles River, France), rats were pair-housed and maintained with *ad libitum* access to food and water under a reversed light cycle (12 hours dark/light cycle; lights off at 10 AM) in controlled temperature conditions, and with a transparent red tunnel as environmental enrichment (8 cm diameter, Bio-Serv, # K3325). Animals were left undisturbed in their home-cages for approximately three weeks, allowing rats to habituate to our Vivarium Facility and routines, and to reverse their circadian rhythm. After this period, animals were handled four times every other day during one week. Body weight was controlled weekly and prior to each testing session. Experiments were performed during the dark cycle, waiting at least 2 hours after the lights were off to start with behavioral procedures. Animals were provided by a commercial company, thus previous social experience, social status and degree of relatedness between the animals was not known. Animal husbandry and all experimental procedures were approved by the Animal Care and Users Committee of the Champalimaud Neuroscience Program and the Portuguese National Authority for Animal Health (Direcçao Geral de Veterinaria), which is in strict compliance with the European Directive 86/6097EEC of the European Council. We confirm that the study is reported in accordance with ARRIVE guidelines (https://arriveguidelines.org/).

### Experimental Procedures

Twenty pairs of animals were tested in the different behavioral paradigms (Sucrose Competition, Food Competition with and without food deprivation, Tube test and Water Competition, see below for description of each task) in order to identify social rank and study reliability of social status between them. Each pair of animals consisted of cage mates living in the same cage for 4 weeks before starting the behavioral procedures. The interacting animals were thus familiar and the same pairs were maintained throughout the entire duration of the experiment. In order to control for possible influences of the order of behavioral testing on the evaluation of social status, we divided the animals in two independent groups (n=10 pairs each) where the order of the tests that involved competition for positive reinforcers was counterbalanced (Supplemental Figure 1). Those tests that required food or water deprivation, and thus were more stressful and/or could a priori induce strong aggressive behaviors, were performed towards the end of the experiment. All pairs were tested in all the tasks with a 2 to 6 days of interval between testing sessions.

At the beginning of the experiment, we randomly identified each rat of a cage as ‘Animal A’ and ‘Animal B’ and quantified their behavior and consumption of the resources for each of the behavioral tests. Before each habituation or test session the fur of the animals was marked using a black pen in order to enable clear identification of each animal for post hoc video-annotation analysis. All the tasks, except for the Tube Test, were performed in the animal’s home cage with small modifications to the lid to accommodate a customized feeder/water bottle. During testing, standard chow and water bottles were removed from the home-cage, to accommodate the modified lids, being replaced immediately after behavioural testing.

### Modified Food Competition test (mFC)

Food competition for palatable pellets (Dustless Precision Pellets, 45mg, Rodent Purified Diet, Bio-Serv) was performed in the home-cage of non-food restricted pairs of animals. For this test, the home-cage lid was replaced by a modified laser-cut acrylic one that accommodated a fully transparent feeder (Figure 1A). The feeder was designed so only one animal could access the food pellets at a time, promoting conflict and competition for the reward. Moreover, the feeder accommodated a sliding door that prevented access to food pellets during the inter-trial interval, and an opening on the top to facilitate delivery of food pellets in each trial with minor interference from the experimenter. This customized lid was used during habituation and test sessions.

Animals were exposed to palatable pellets in their home-cages during the handling period for four days in order to reduce neophobic responses to the food. Then, during three consecutive days, all the animals went through a habituation period to the modified lid, where they were allowed to explore and consume the pellets individually without competition, while the partner would be kept in a separate cage. Specifically, during habituation days, the new lid holding the feeder was placed on the home cage containing 10 palatable pellets. The sliding door was closed, preventing access to the pellets. Two minutes after, the door was opened, allowing the rat to access the pellets for 2 minutes, after which the door was closed again and 10 new pellets were placed. In total, the animal was given 4 minutes’ access to 20 pellets in a total session of 10 minutes. Next, food competition in a social context was performed for two consecutive days. Pairs of animals were re-marked, and the home-cage lid was replaced by the modified one with the feeder and 10 palatable pellets. 1 minute after, the sliding door was opened allowing the rats to have access to the pellets for 2 minutes, after which the door was closed again for a 1-minute inter-trial interval and 10 new pellets were delivered. We repeated this procedure for 5 trials and a total session of 15 minutes and 50 pellets. After the session, the customized lid was replaced by their home-cage lid.

### Modified Food Competition with food deprivation (mFCD)

The modified Food Competition test was performed in the home cage of familiar animals as described above, but in this case animals underwent only one session of social competition after a 24h period of food deprivation. After the test, the modified lid was replaced by the standard one and rats were allowed to eat and drink *ad libitum* for the rest of the cycle.

### Sucrose Competition (SC)

Pairs of non-water-restricted cage mates competed for access to a bottle containing 1% sucrose solution placed in a modified lid on their home-cages. The lid was designed so the bottle holders were prolonged with a transparent acrylic tube, in a manner that the tip of the bottle was surrounded by an extension that would allow the head of only one animal to drink at a time (Figure 1B). We performed three habituation sessions. In the first habituation day, animals were exposed to the new lid for 20 min where no bottle was available. In the two following days, the new lid was holding two bottles of 1% sucrose solution and animals were given free access to the sucrose solution for 20 minutes. Then, animals were re-marked and tested for sucrose competition in two consecutive days for 10 minutes, where the modified lid presented only one bottle of sucrose this time.

### Sucrose Competition with intermittent access to the resources (SCI)

In this test, access to the 1% sucrose solution followed an intermittent schedule. One minute after the beginning of the session, a bottle with sucrose solution was placed in the dispenser allowing the animals to access the solution for 2 minutes. After this time, the bottle was removed for 1 minute and put back again for 2 more minutes, performing a total of 5 trials. The SCI was performed over two consecutive days. After each session, the customized lid was replaced by their home-cage lid.

### Water Competition (WC)

Animals were water deprived for 24h and tested for competition for water in their home cage. At the moment of the test, the cage lid was replaced by a modified one where access to the bottle was only possible for one animal at a time. The duration of the test was 10 minutes, after which the standard lid was replaced and both rats had *ad libitum* water access.

### Tube Test (TT)

We used a transparent Plexiglas tube with 60 cm length and 8 cm diameter, a size that allows an adult rat to pass through without reversing its direction, and, when two rats are placed in the tube, prevents one rat from crossing the tube by passing the other. We performed one habituation session where animals individually explored the apparatus, allowing spontaneous entering and crossing of the tube for 5 minutes. During this habituation session, animals were placed initially in front of one of the ends of the tube, and freely allowed to enter the tube and explore the behavioural table. In our hands, rats immediately entered and crossed the tube, spontaneously performing 5 to 6 crossings during the habituation period, without the need to force exploration nor to push them, as they did not display extended periods of immobility. After the 5 minutes’ habituation animals were returned to their home-cages. During test days, each pair of cage mates rats was simultaneously placed into opposite ends of the tube and met in the middle. At this time, a partition placed at the center of the tube was removed. The rat that first retreated from the tube was designated as the ‘loser’ and the other as the ‘winner’. After each trial, both rats were placed back into their home-cages until the beginning of the next trial. From trial to trial, animals were released at either end of the tube alternately. We performed one testing session of the Tube Test with 10 trials at the beginning of the experiment and assessed stability of social rank within this test with another testing session at the end of the experiment. We quantified the amount of time that one of the animals took to push the other out of the tube and annotated the winner and loser of each interaction.

#### Video acquisition and Behavioral Quantification

Experiments were performed under the dark cycle of the animals and video recordings were obtained by a high resolution infra-red camera (PointGrey Flea3-U3-13S2M CS, Canada) under infra-red illumination, capturing frames at 30Hz at 1280×960 pixel resolution. Supervised offline frame by frame video annotation of behaviors of interest was performed by a trained blind experimenter (DFC) after confirmation of highly reliable quantifications. For each animal of the dyad we quantified frequency, latency, and duration of (1) consumption of resources (number of pellets eaten or time spent drinking water or sucrose); (2) exploration of the feeder (sniffing behavior inside and outside the feeders or bottle holders); (3) self-grooming; and (4) pushing the other animal to gain access to the resource. In those tests with a trial structure, behaviors were aligned to time 0, i.e. the moment where the sliding door that gave access to food pellets was opened for food competition tests, or when the bottle was placed in the modified lid in the case of the Sucrose Competition Intermittent. For these tests with trial structure, intertrial interval (the time where reward was not present) was set to 60 seconds and the duration of a trial (where animals could access to the reward) to 120 seconds. However, due to the manual control of the timings of the experiment, some trials of some pairs of animals resulted with shorter durations than aimed. In order to ensure comparable behavioral profiles across trials and dyads across these tests, we decided to narrow our behavioral quantification window and focus our statistical analysis to the 40 seconds before and 80 seconds after time 0 (the moment where access to rewards was possible). This resulted in a total of 10 min duration test for both tasks with and without trial structure. However, note that consumption in tasks with trial structure was only quantified for 400 seconds (80 seconds during 5 trials in a day), while in the other tasks this could be possible during 600 seconds. To take this difference into account, comparisons between consumption levels across tasks were performed with the percentage of time spent consuming relative to the duration of the session. Exploration of the feeder was measured during the whole session, and in those tests with a trial structure, we differentiated between exploration of feeder when reward was accessible and before that, when the sliding door was closed or there was no bottle there yet, as a proxy for anticipatory exploration. Pushing behavior was divided in two distinct categories depending on the outcome: Successful Pushing, if the animal managed to displace the partner from having access to the resource and Unsuccessful Pushing, when animals would attempt to get access to the resources but were unsuccessful to displace their partner from the reward area. In the Tube Test, we quantified the number of wins for each animal and the duration of each trial as a proxy for the time the animals used to solve the territorial conflict.

Bonsai ^49^ and Python Video Annotator (https://pypi.org/project/Python-video-annotator/), both open source computer vision software available online, were used to perform behavioral quantification. First, digitally assigned behaviors were quantified with Bonsai, which created timestamps for the beginning and ending of each behavioral event. Then, the start and end of each behavioral bout was curated with frame-by-frame investigation using Python Video Annotator, which allows fine modification of the timeframes with subsecond resolution. Moreover, Python Video Annotator allowed easy post hoc categorisation of the two types of exploration of the feeder (anticipatory or during the presence of the resource) and pushing behavior (successful or unsuccessful) which can only be identified once the bouts of pushing behaviour are finished and is thus not possible to analyse with online video analysis.

### Data Analysis

Data was parsed and processed with Python (Python Software Foundation, v.2.7). In addition to comparing raw data obtained, we calculated several indexes to compare hierarchies across tests.

#### Dominance Index

The *Dominance Index* (DI) was calculated for each test, where the difference in resource consumption across partner animals was normalized by the summed resource consumption of the pair, following this formula:

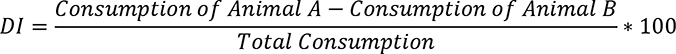

Consumption corresponded to the number of pellets in the modified Food Competition tests, the duration of drinking in the Sucrose and Water Competition tests, and the number of wins displayed by an animal in the case of the Tube Test. The sign of this index would indicate whether animal A or B would consume more, i.e. positive values would indicate that animal A consumed more, and negative values that animal B consumed more. DIs close to 0 would indicate no differences in consumption between the animals of a pair. Differences of 5% to equal consumption, i.e. DI ranging from −10 and +10, were considered noise and indicative of no reliable hierarchy.

#### Conflict Resolution Index in the Tube Test

The *Conflict Resolution Index (CRI)* was calculated taking into account not only who won a trial, but also how long it took for the conflict to be resolved (i.e., the latency for one of the animals to be pushed out of the tube):

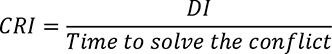

#### Conflict Index

The *Conflict Index* (CI) was calculated by dividing the time animals spent pushing in a specific test by either: (1) the time spent consuming, in case of the Sucrose and Water Competitions, or (2) the latency to eat all rewards in the case of the modified Food Competition with and without food deprivation.

#### Statistical analysis

The statistical analysis was performed using IBM SPSS Statistics version 24.0 for Windows. The normality of the data was tested using Kolmogorov-Smirnov normality test, and when normality was not observed non-parametric tests were applied and median and 95% confidence interval were chosen to represent data in figures. Wilcoxon signed-rank tests with Bonferroni correction were used to study differences between counterbalanced groups in each protocol, of each behavior across the tasks, and to study differences between dominant and submissive animals on behavior. Paired t-test were performed to assess differences in the weights between dominant and submissive animals. One-Way ANOVA followed by post-hoc test Tukey was used to compare behaviors of interest across tasks. Here, when normality was not observed, a Kruskal-Wallis test with post-hoc Dunn-Bonferroni correction was performed. Bivariate Pearson Correlation was performed to measure the strength and direction of association between the Dominance Index of all the tasks and linear regressions for assessing predictive value of Food Competition and Tube Tests controlling for body weight. Statistical significance was set at p<0.05.

## Acknowledgements

This work was supported by grants of the NARSAD Young Investigator Grant from the Brain & Behavior Research Foundation under the grant number 26478 to C.M., the Spanish Agency of Research (grant RTI2018-097843-B-100 to C.M.), the “Severo Ochoa” Program for Centers of Excellence in R&D (SEV-2013-0317 and SEV-2017-0723) and the Champalimaud Foundation. D.F.C. was further supported by the Ministerio de Ciencia e Innovación (BES-2016-07674) and C.M. by a Ramon y Cajal contract (RYC-2014-16450).

We thank the Marquez lab for fruitful discussions and specially, Kevin Caref for insightful comments on the manuscript. We also express our gratitude to Gonçalo Lopes for his help with Bonsai workflow for annotation of behavior, Antonio Dias for feedback on the python scripts and to Cristina Savin for discussion of the data analysis.

## Author contributions statements

**DF Costa**: Conceptualization, Investigation, Methodology, Formal analysis, Visualization, Writing original draft and Review and editing the manuscript; **MA Moita**: Methodology, Review and editing manuscript, Funding acquisition; **C Márquez**: Conceptualization, Supervision, Investigation, Methodology, Formal analysis, Visualization, Writing - original draft, Writing - review & editing, Funding acquisition.

## Additional information

### Data availability

All data generated to support the findings of this study are available from the corresponding author upon reasonable request.

## Supplemental Figures

**Supplemental Figure 1:**
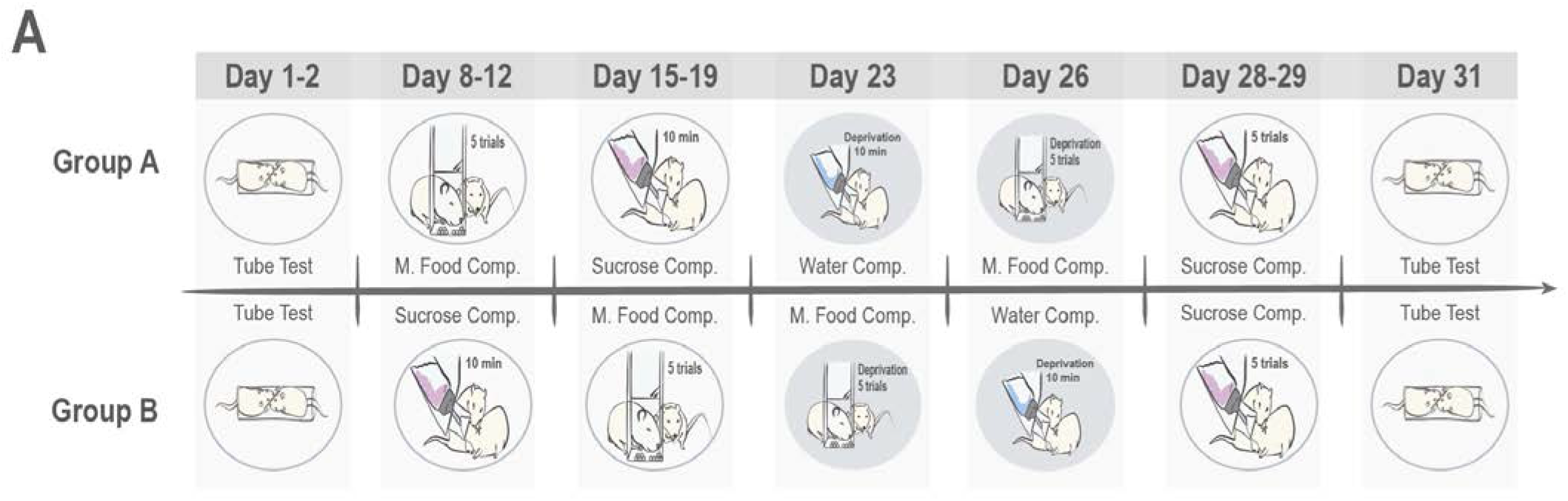
Timeline of experimental design. After handling, two independent groups of animals were created where the order of the tests was counterbalanced. The order of the test is provided in the cartoon, and days indicate when they were performed. Note that in the case of the first evaluation of the Tube test, the modified Food Competition test without deprivation and the Sucrose Competition with continuous access to the bottle (10 minutes), these days include also the habituation sessions. All animals were tested in all competition tasks with inter-tasks intervals ranging between 2 and 6 days. Sucrose competition with intermittent access to the bottle (5 trials) was included towards the end of the experiment, after realization of the low amounts of drinking performed in the sucrose test with the continuous access to the bottle configuration. Shaded circles in the competition tasks at days 23 and 26 indicate that they were performed under deprivation states.

**Supplemental Figure 2:**
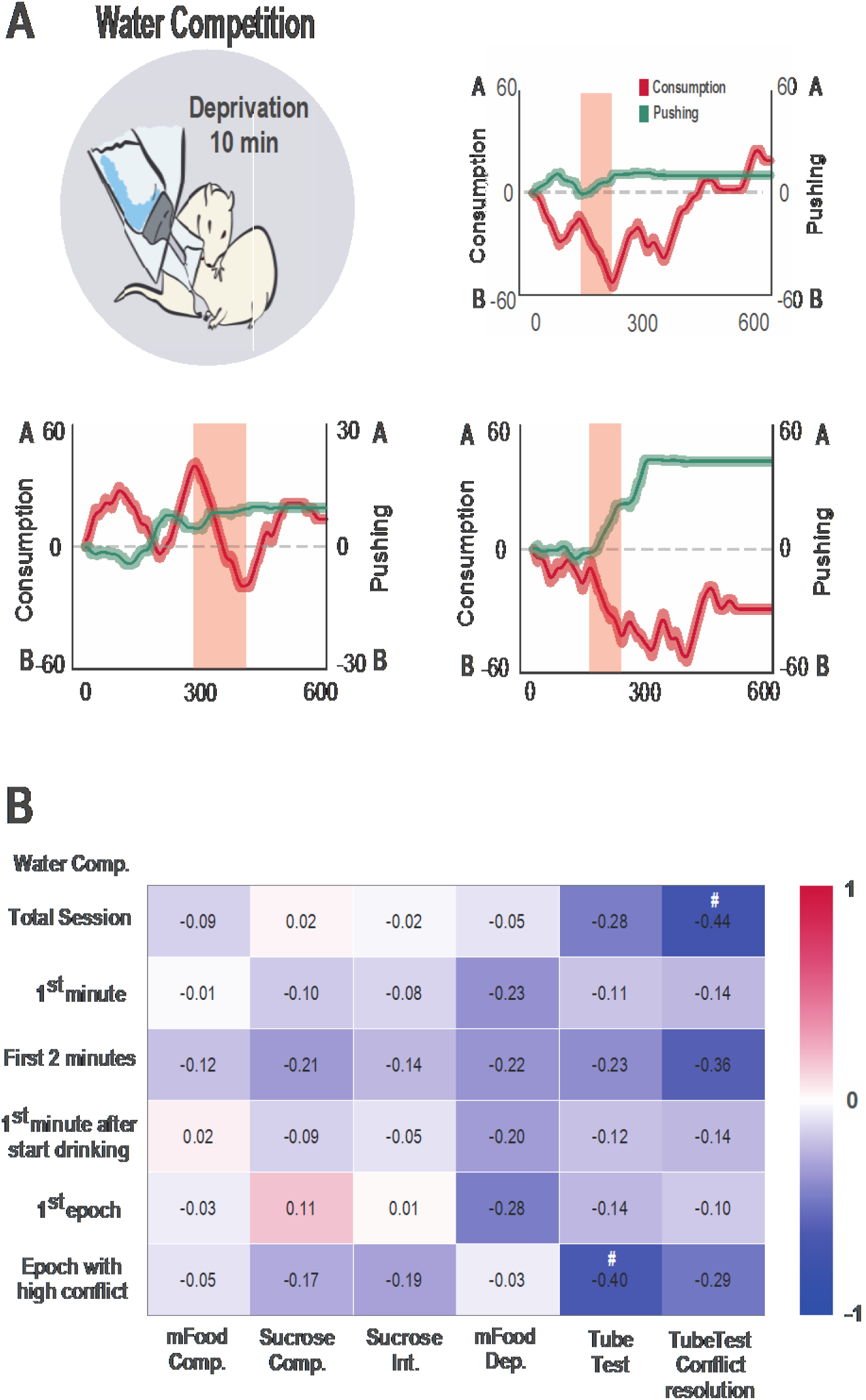
Identification of moments with highest conflict in the Water Competition tests and lack of correlation with Dominance Index (DI) in any other test. **(A)** Graphical representation of dynamics of consumption (red, left axis) and pushing behaviors (green, right axis) and selection of the bout with highest conflict (shaded red rectangle) in three example pairs of animals during Water Competition test. Drinking and pushing raw data was transformed into modified cumulative plots and smoothed using convolution with a Gaussian filter of 30ms standard deviation. In this plots the direction of the cumulative graph indicated which animal was drinking or pushing over the session, either animal A or B. In this way, increases in this cumulative plot would indicate that animal A would be drinking, decreases that animal B would drink, and flat stable lines that no animal was drinking. The same applied for pushing data (in green). For example, for the example pair 1 animal B would start drinking while animal A was pushing for around 100 seconds, then they would alternate for a brief period of time, followed by another alternation, where a long bout of drinking was performed again by animal B while animal A continued pushing. After that, no significant pushing was performed by any of the animals, and although some alternations in drinking would be observed, now animal A would take over and drink more. In the different example graphs we can observe that dynamics between pairs are different over time but that animals mostly alternate in their drinking times, and pushing behavior decreases around half of the session. In order to select the bouts of highest conflict, we first defined epochs of consumption and pushing displayed by the pair by identifying the turning points that marks the moments when significant changes in the behavior occurs (x[n + 1] − x[n] < 0). For each epoch we calculated the duration of both consumption and pushing behaviors and selected the epoch with the highest value of both consumption and time pushing the partner from the water dispenser. (B) No significant correlation was observed between Dominance Indexes (DI) of Water Competition when calculated with the drinking duration in the whole session, nor in the first min or the two first minutes, nor when taking into account the moment when animals started drinking. We then identified the epochs with longest drinking, as they could be not necessarily in the early moments of the test, but no correlation was observed with other behavioral tasks either. We then selected the epochs with higher conflict (see A for representative examples of the identification of the epochs), but again, no correlation was observed. Values in the correlation matrix correspond to Pearson correlation r values. #p<0.10.

**Supplemental Figure 3:**
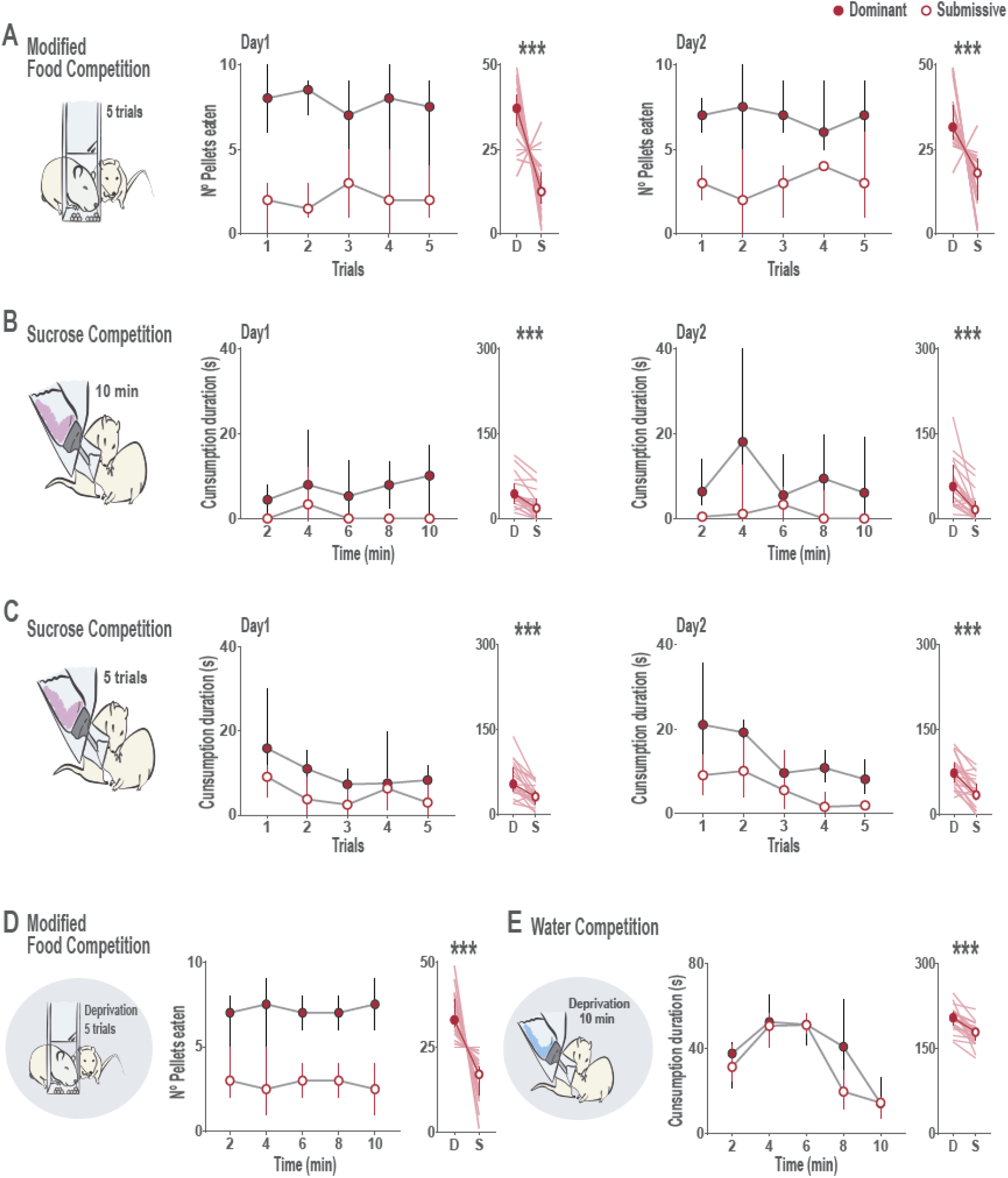
Social hierarchy measures are stable across time. Consumption of resources is plotted over time or trials within a session and across days. In all cases the total consumption of all testing days was taken as the criteria to define dominant and submissive animals. In those tasks with a trial structure, results for each day are first presented by trials and then as the average of consumption in that day. For the sucrose competition with continuous access and the water competition, results are first presented in 2 min blocks and then the average consumption of that day. Statistics evaluating differences between dominant and submissive animals were performed in the average consumption of a day (**A**) consumption in the modified Food Competition on each of the two days of testing when the criteria to define hierarchy was the total consumption of the two days. Dominant and submissive animals clearly differed in their consumption across time. (**B**) Similar for the Sucrose Competition with continuous access to the bottle. Note the low levels of consumption, especially in day 1. (**C**) Similar for the Sucrose Competition with intermittent access, (**D**) the modified Food Competition under deprivation and (**E**) the Water Competition. Median, 95% CI and individual values for all animals are represented. ***p<0.001 after non parametric Wilcoxon test.

**Supplemental Figure 4:**
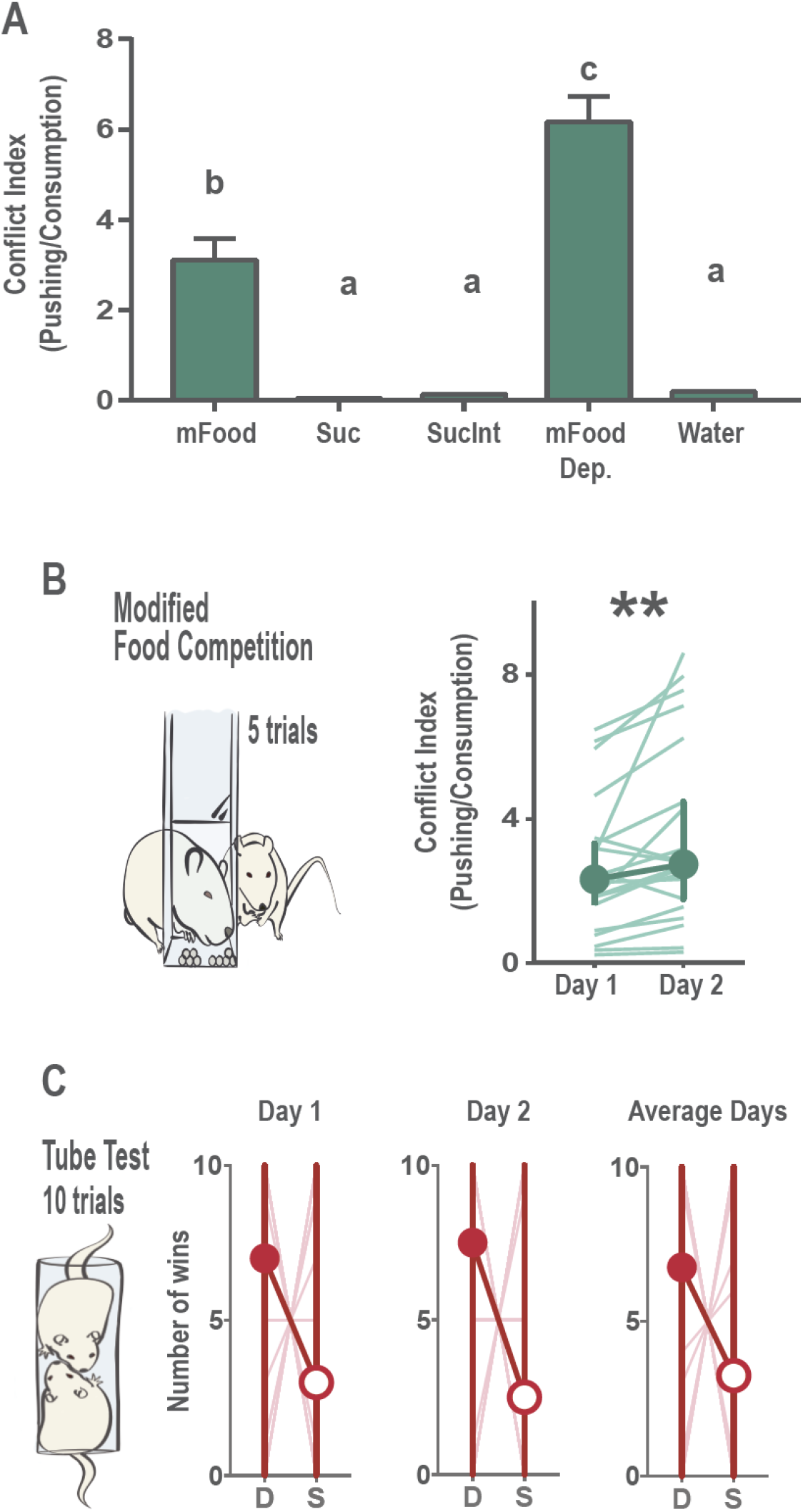
Food Competition tests reflect higher conflict. (**A**) We calculated a conflict index for each test that involved consumption of resources by dividing the duration of pushing performed by the time the spent consuming (as the duration to consume 1 pellet would be few milliseconds in the case of the Food Competition tests, we considered the latency to eat the 10 pellets available in each trial). Due to the short duration of pellet availability and high pushing levels observed in the Food Competition tests, these tests showed higher conflict indexes than the sucrose tests or water competition (F(4, 98)=64.805 p<0.0001 followed by Tukey posthoc), which was more marked when animals were under deprivation. Average and SEM are represented, and letters denote statistically significant differences between behavioral tests after one-way ANOVA with Tukey posthoc comparisons. (**B**) Conflict index was higher in the second day of testing in the modified Food Competition test (paired t-test t(19)=-2.985 p=0.008). Median, 95% CI and individual values for all animals are represented. ** p>0.01. (**C**) Dominant animals defined according to their behavior in the second day of modified Food Competition test did not differ in the amount of winnings in the tube test in either of the days tested nor when the average winnings of the two days were considered (Wilcoxon signed-ranks for TT Day1: z=-0.774, p=0,44; TT Day2: z=-0.503, p=0,62; TT AVG Days = z=-0.699, p=0,49). Median, 95% CI and individual values for all animals are represented

